# Transforming avian to mammalian modes of digit patterning

**DOI:** 10.64898/2026.06.30.735497

**Authors:** Sarah Wiggins, Sofia Sedas Perez, Marysia Placzek, Rory Cooper, Matthew Towers

## Abstract

How a conserved embryonic organiser—the Sonic hedgehog (Shh)–expressing Zone of Polarising Activity (ZPA)—generates diverse limb architectures in amniotes remains a central problem in evolutionary and developmental biology. The principal difference across species lies in the number of digits produced from ZPA tissue: two in mammals^1^, one in the chick leg^2^ and none in the chick wing^2^. Here we show that this divergence is governed by a Shh–p27^Kip1^ pathway operating in avian, but not mammalian ZPAs. In chick wing ZPA explants, attenuation of this pathway reveals an intrinsic digit-forming programme, enabling cells to self-organise signalling networks and generate digits after grafting into a host wing bud. Guided by these findings, we redirected the chick leg, which retains stem amniote digit identities, to follow a mammalian-like developmental trajectory. Precise temporal restriction of Shh signalling transforms chick legs into pentadactyl limbs with digit identities characteristic of mammals and their therapsid ancestors, with two digits arising from the ZPA. These findings establish a unifying framework for how Shh controls both digit number and identity across amniotes.

## Main

The digits of the vertebrate limb provide a classic system for studying how developmental mechanisms shape morphological and functional diversity. The fossil record indicates that the ancestral pentadactyl (five-digit) pattern of stem amniotes (Fig. 1) was reduced to three digits with fewer joints (phalanges) in the avian wing^3,4,5^, a transition that facilitated the evolution of flight (Fig. 1 - pf, phalangeal formula; FL, forelimb). By contrast, the avian leg was reduced to four digits, which retained the evolutionary ancient stem amniote identities defined by a conserved phalangeal formula (Fig. 1 - pf 2-3-4-5; HL, hindlimb)^3,4,5^. Mammalian forelimbs and hindlimbs typically retained the pentadactyl digit pattern, but with reduced numbers of phalanges, a modification associated with the shift from a sprawling to an upright gait^6,3^ (Fig. 1).

**Figure 1:**
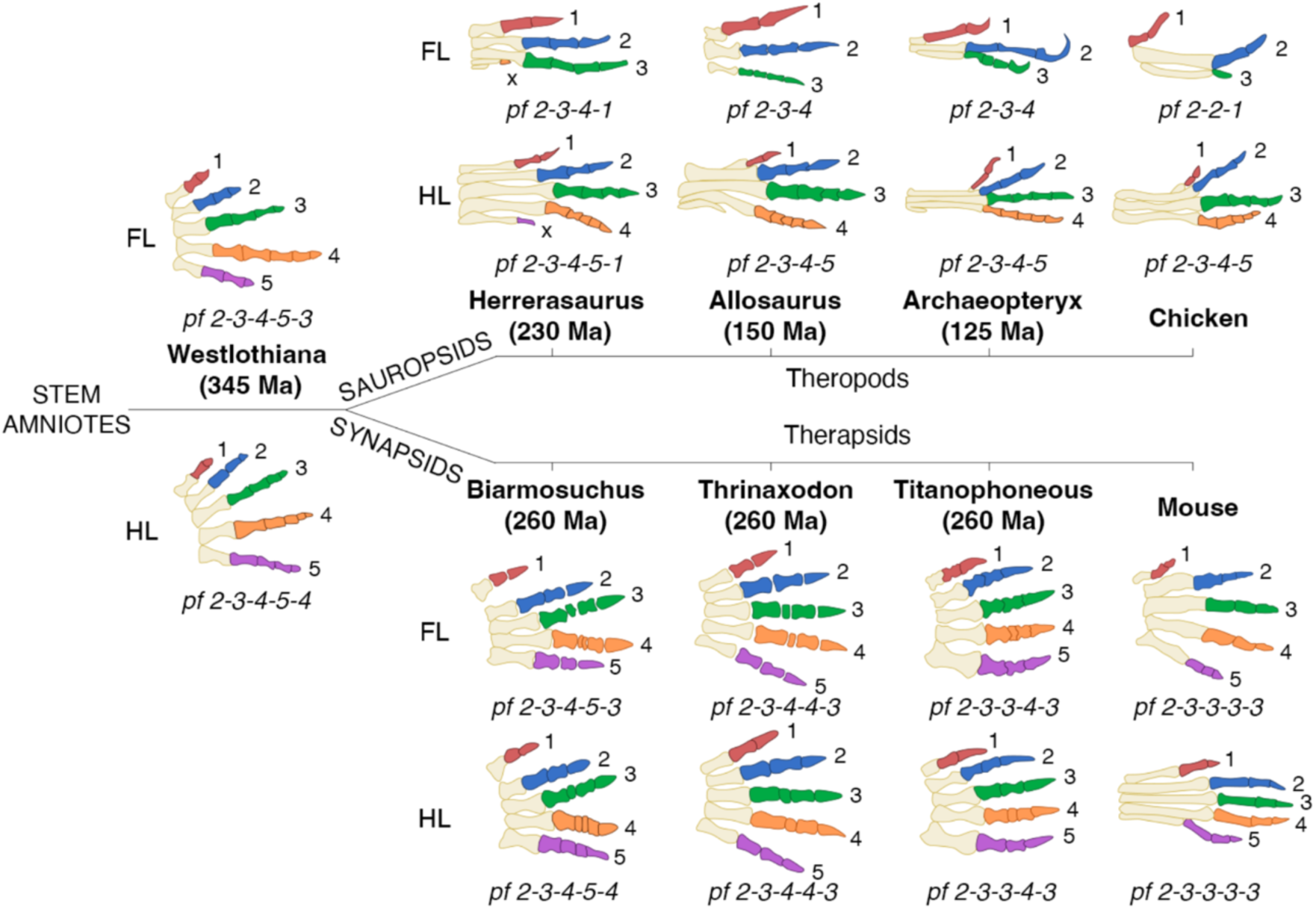
Avian and mammalian digit evolution. **a**, Schematic comparison of amniote digit patterns and phalangeal formulae (pf). The ancestral pentadactyl (five-digit) stem amniote forelimb exhibited a pf of 2-3-4-5-3 (anterior-to-posterior) and 2-3-4-5-4 in the hindlimb (digit identities are colour-coded)^19^. In the avian lineage, wing digits were reduced to a pf of 2-2-1 following the loss of digits 4 and 5. Avian legs retain the ancestral pf (2-3-4-5) but have lost digit 4^19^. Mammalian limbs generally maintain pentadactyly and exhibit a reduced pf of 2-3-3-3-3^6^. a, anterior; p, posterior; pr, proximal; d, distal.

Fate mapping experiments demonstrate that digit number depends on the developmental potential of the Zone of Polarising Activity (ZPA), a mesodermal organiser located at the posterior margin of the limb bud that produces Sonic hedgehog^7^ (Shh). The ZPA produces no digits in the chick wing, one in the chick leg and two in mouse limbs^1,2^. Subsequent studies in the chick wing showed that—in addition to specifying digit identity in tissue adjacent to the ZPA by a morphogen gradient—Shh activates *p27^Kip1^*, a G1/S phase D-cyclin-dependent kinase inhibitor, thereby limiting proliferation in the ZPA^8,9^. Experimentally, precise temporal inhibition of Shh signalling causes ZPA cells to lose *p27^Kip1^* expression, over-proliferate and form an additional ZPA-derived posterior digit^8,9^—a process that also requires signalling from the overlying Apical Ectodermal Ridge (AER)—a distal epithelial thickening that expresses *Fgf8*^8^.

Although two digits arise from the mouse limb ZPA^1^, genetic studies indicate that digit patterning requires only a transient pulse of Shh signalling, independent of a morphogen gradient^10^. How this conserved signalling system generates the distinct digit numbers and identities of birds and mammals therefore remains unresolved, creating a significant disconnect between classical chick embryology, mouse genetics and the fossil record. Moreover, comparative transcriptomic analyses indicate that, apart from the thumb, digit identities cannot be reliably assigned across amniotes using gene expression profiles^11^, limiting the utility of this approach for understanding how divergent digit patterns evolve. The intricate crosstalk between the ZPA and AER has also obscured whether digit-forming ability is intrinsic to the ZPA itself and how this capacity has been modified during evolution. To address this, we developed a chick wing ZPA explant system that enables its developmental potential to be examined independently of the AER.

### The ZPA possesses an intrinsically timed programme

To establish an ex vivo chick wing bud ZPA explant (ZPAE) system, we adapted our existing methodology for culturing distal mesoderm^12^. The ZPA was excised from wing buds at stage HH19-20 (Hamburger Hamilton 19-20), when S*hh* expression is established, and this time-point was designated as 0 h (Fig. 2a). To verify the accuracy of ZPA excision, we performed fluorescent multiplex Hybridisation Chain Reaction (HCR) in situs, which confirmed that *Shh* is consistently expressed throughout the excised ZPAs, while neither AER-restricted *Fgf8*, nor mesodermal *Grem1*—which encodes the AER maintenance factor^13,14^—are detectable (Fig. 2b; Supplementary Figure 1). Cultured ZPAEs faithfully maintain in vivo timing of *Shh* expression, which persists for approximately 48 h^15^, demonstrating that its duration is intrinsically determined, and is independent of extrinsic signalling between the distal mesoderm and the AER (Fig. 2c-f). Over time, *Shh* expression becomes restricted to distal regions of the ZPAEs, while non-*Shh*-expressing descendants are displaced proximally by growth (Fig. 2c-f). Thus, ZPAEs maintain key in vivo developmental parameters in the absence of positional cues from adjacent mesoderm or an overlying AER.

**Figure 2:**
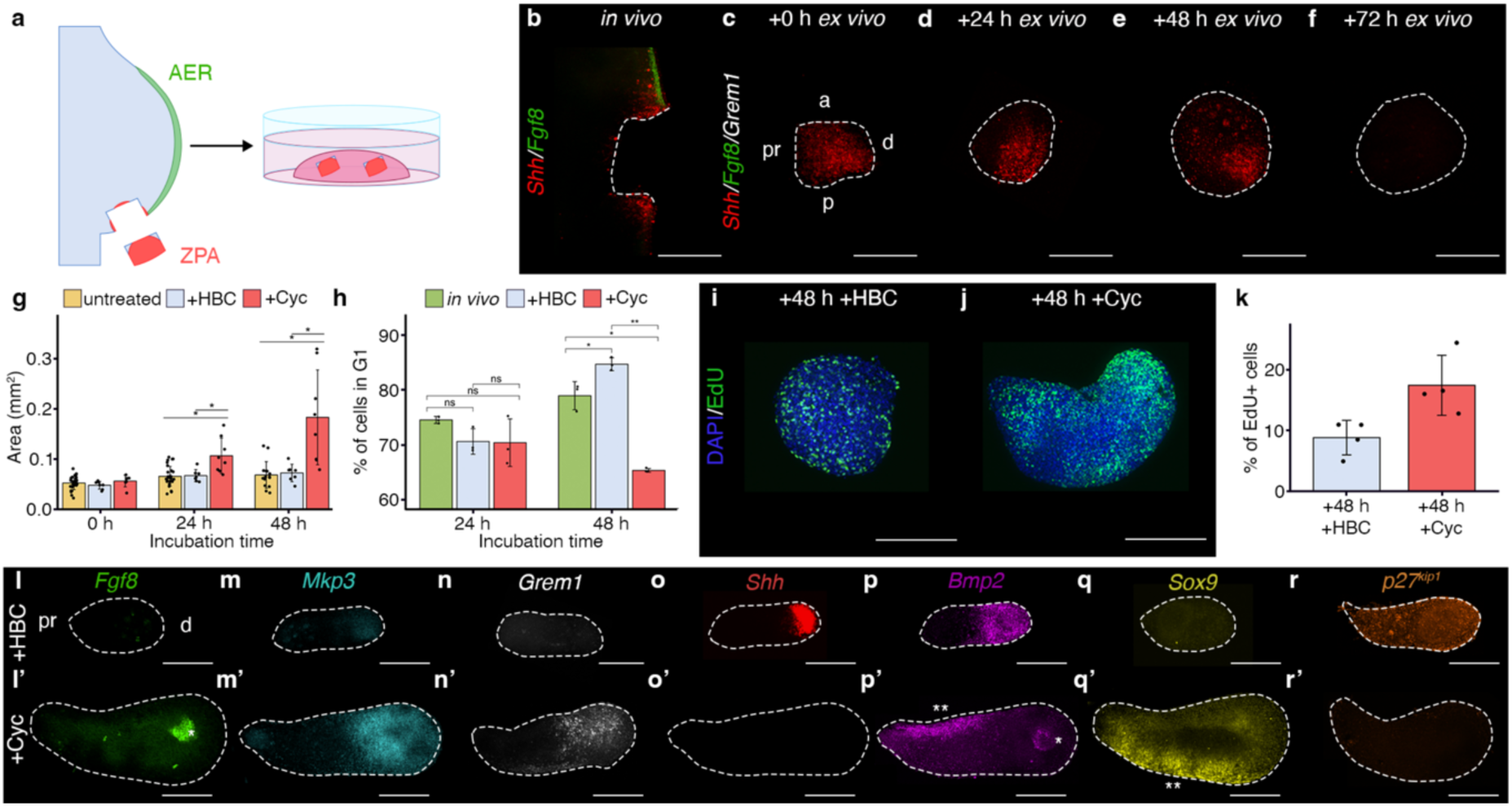
Chick wing ZPAEs maintain intrinsic developmental parameters and respond to Shh inhibition. **a**, Schematic of the procedure to isolate the ZPA (red) from surrounding mesenchyme (blue) and AER (green) at HH19-20 for Matrigel culture. **b, c**, Multiplex HCR RNA-FISH of freshly isolated ZPA explants (ZPAEs) at 0 h. *Shh* is expressed throughout (n = 13/13), while *Fgf8* (n = 10/13) and *Grem1* (n = 13/13) are absent. **d–f**, HCR RNA-FISH analysis showing that *Shh* expression is maintained for 48 h (24 h: n = 42/42; 48 h: n = 15/15) and is undetectable at 72 h (n = 12/12). **g**, Quantification of ZPAE surface area. Inhibition of Shh signalling via cyclopamine (cyc) results in significant overgrowth compared to carrier-controls (HBC) (n = 8, 7; *P* = 0.016) and untreated ZPAEs (n = 8, 29; *P* = 0.013) at 24 h and 48 h (n = 8, 7; *P* = 0.013 and n = 8, 15; *P* = 0.011 respectively); no significant difference was observed at dissection (n = 7, 7; *P* = 0.146). **h**, Flow cytometric analysis of G1-phase cells revealed no significant difference at 24 h between cyc-treated ZPAEs and both carrier-controls (n = 3, 3; *P* = 0.942) and in vivo ZPA cells (n = 3, 3; *P* = 0.474). At 48 h, a significant difference was observed between cyc-treated ZPAEs and both controls (HBC; n = 3, 3; *P* = 0.002; in vivo; n = 3, 3; *P* = 0.021). **i–k**, EdU labelling (**i, j**) and quantification (**k**) at 48 h. Cyclopamine treatment significantly increases the proportion of S-phase cells in distal regions compared to HBC (n = 4, 4; *P* = 0.0311). **l–r**, Multiplex HCR RNA-FISH analysis at 48 h comparing HBC and cyclopamine-treated ZPAEs. **l, l’**, *Fgf8* is induced de novo in the distal (d) end of cyc-treated explants (n *=* 5/8) but remains absent in controls (n = 7/9). **m, m*’***, Distal up-regulation of the FGF-target *Mkp3* following Shh inhibition (n = 4/4) compared to weak basal expression in controls (n = 3/3). **n, n’**, Ectopic distal expression of *Grem1* in cyc-treated explants (n = 2/3). **o, o’**, Loss of *Shh* expression following cyclopamine treatment (n = 14/15). **p, p’**, Re-localisation of *Bmp2* in cyc-treated explants to distal (*) regions and a proximal anterior (**) stripe (n = 4/4). **q, q**’, Expression of *Sox9* in a proximal posterior stripe (**) following Shh inhibition (n = 5/5). **r, r’**, Loss of *p27^Kip1^*expression in cyc-treated ZPAEs (n = 4/5) compared to uniform expression in controls (n = 4/4). a, anterior; p, posterior; pr, proximal; d, distal. Statistical significance was determined via two-tailed unpaired t-tests. Scale bar, 250 µm.

### Shh signalling restrains ZPA proliferation and growth

To determine how Shh signalling regulates ZPAE development, we inhibited pathway activity at the level of Smoothened by adding cyclopamine to the culture medium at 0 h. The loss of *Ptch1*—a downstream transcriptional target of Shh—confirmed effective pathway attenuation (Supplementary Fig. 2). Quantification of explant surface area revealed that control ZPAEs (untreated or HBC carrier–treated) expanded by 1.30- and 1.50-fold, respectively, over 48 h, with most growth occurring during the first 24 h (Fig. 2g). By contrast, cyclopamine-treated ZPAEs expanded by 3.22-fold over the same period and typically developed a broader distal than proximal domain (Fig. 2g; Supplementary Fig. 3).

Flow cytometric analysis showed that control ZPAEs maintain the in vivo decline in proliferation rates over 48 h, as reflected by increased proportions of cells in G1-phase (Fig. 2h). Inhibition of Shh signalling did not alter proliferation at 24 h; however, by 48 h the proportion of G1-phase cells decreased by 19.28%, consistent with accelerated G1-to-S phase progression (Fig. 2h). Enhanced proliferation was verified by EdU incorporation, revealing a 1.97-fold increase in S-phase cells at 48 h, predominantly localised to distal regions of the explants (Fig. 2i-k). Together, these results demonstrate that Shh signalling restrains ZPAE growth by slowing progression into S-phase.

### Shh signalling prevents the ZPA from organising local signalling networks

To investigate how Shh signalling regulates ZPAE behaviour, we examined gene expression at 48 h using HCR multiplex in situ hybridisation. In controls, *Fgf8* was undetectable, whereas cyclopamine-treated ZPAEs exhibited strong distal expression (Fig. 2l), accompanied by a significant up-regulation of its downstream transcriptional target, *Mkp3*, in adjacent regions (Fig. 2m). Similarly, *Grem1* was undetectable in controls but up-regulated in distal regions of cyclopamine-treated ZPAEs (Fig. 2n). In controls, *Shh* and *Bmp2* were co-expressed in overlapping distal domains, recapitulating in vivo patterns^16^, which were lost following cyclopamine treatment (Fig. 2o–p). Instead, *Bmp2* co-localised with *Fgf8*—resembling in vivo AER expression^16^—and was additionally detected in a discrete anterior stripe (Fig. 2p’; compare with Fig. 2l’). Stripes of *Bmp2* mark the interdigits and induce adjacent *Sox9*-expressing digit condensations through self-organisation, the number of which scales with limb bud width^17,18^. Accordingly, *Sox9* was absent in controls but appeared as a posterior stripe in enlarged cyclopamine-treated ZPAEs (Fig. 2q). In addition, *p27^Kip1^*, which was uniformly expressed in controls, was undetectable in cyclopamine-treated ZPAEs, potentially contributing to their overgrowth (Fig. 2r). Collectively, these findings indicate that Shh signalling prevents ZPA cells from organising signalling networks and initiating chondrogenesis.

### Shh signalling prevents the ZPA from forming skeletal elements

Given that inhibition of Shh signalling can induce digit formation from the chick wing ZPA^8^, we investigated whether pre-chondrogenic cells in cyclopamine-treated ZPAEs can form skeletal elements. 24 h GFP-expressing explants (HH24 equivalent) were grafted to the posterior margin of HH21 wing buds, replacing the host ZPA (Fig. 3a). Control ZPAE grafts contributed to the posterior margin of digit 3 (Fig. 3b; Supplementary Table 1). By contrast, cyclopamine-treated ZPAE grafts produced cartilaginous elements reflecting their proximo–distal position: distal grafts generated one or two segmented digit-like structures (d*), whereas proximal grafts formed a single element resembling the ulna (u*) (Fig. 3c–d; Supplementary Table 1).

**Figure 3:**
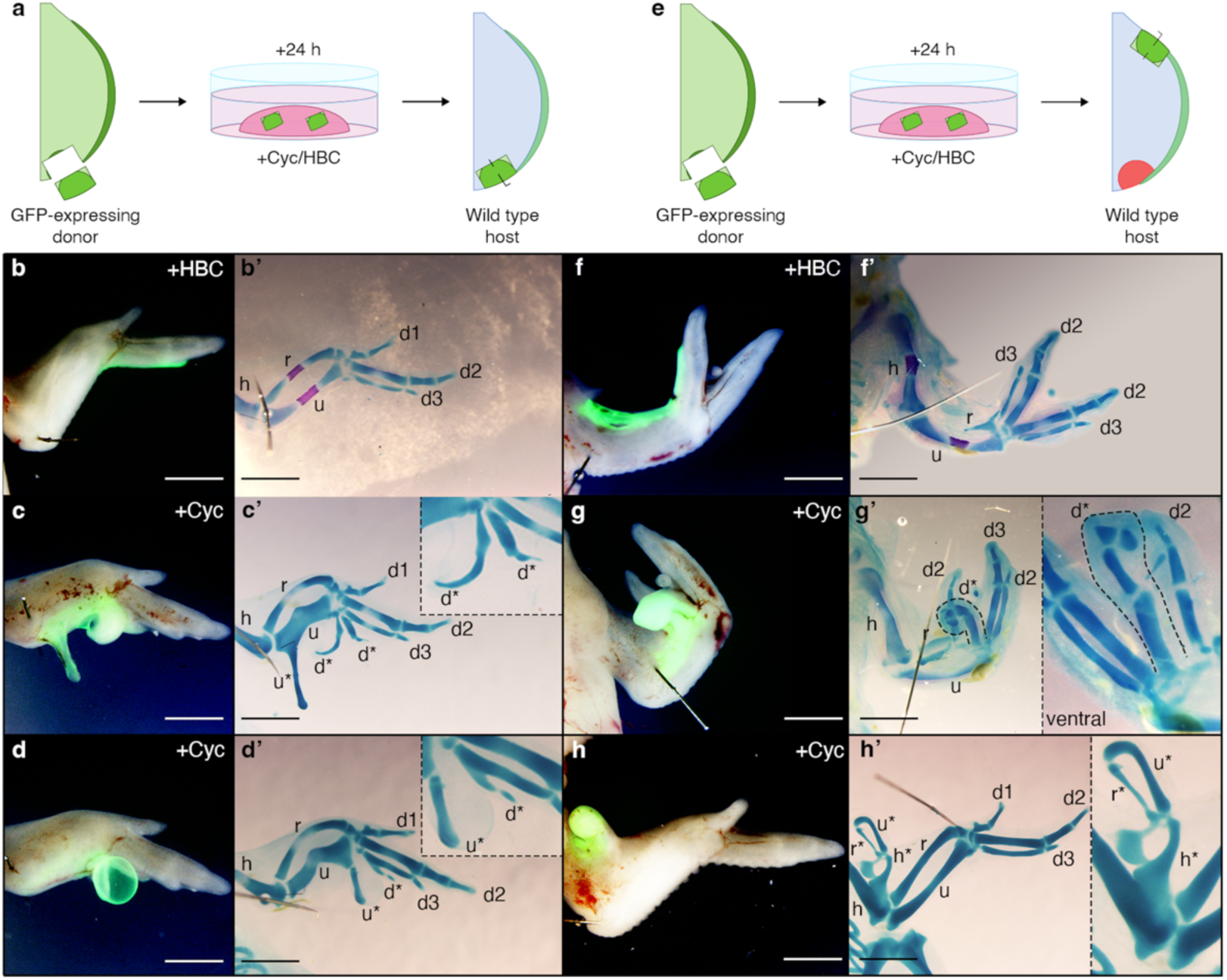
Shh signalling represses the skeletal element forming potential of the ZPA. **a**, Schematic of the grafting procedure. GFP-expressing ZPAEs (HH24 equivalent) were isolated from donor embryos and grafted posteriorly into wild type wing buds in place of the endogenous ZPA at HH21–22. **b–d**, Representative results of grafts placed at the posterior margin. **b, b’**, Control (HBC-treated) ZPAEs contribute to soft tissue flanking the posterior margin of digit 3 without generating additional cartilage (n = 6/7). **c-d**, Cyclopamine-treated ZPAEs produce skeletal elements (n = 3/4). The morphology of elements reflects their proximo–distal position: those projecting from distal regions resemble digits (d*), whereas those projecting from more-proximal regions resemble the ulna (u*) **e**, Schematic of the grafting procedure. GFP-expressing ZPAEs (HH24 equivalent) were isolated from donor embryos and grafted anteriorly into wild type wing buds at HH21–22. **f–h**, Representative results of grafts placed at the anterior margin. **f, f’**, Control ZPAEs typically fail to develop skeletal structures (n = 6/7) and often duplicate the host pattern of digits. **g–h**, Cyclopamine-treated ZPAEs produce skeletal elements (n = 7/10). Elements projecting from distal regions resemble digits (d*) and often duplicate the host pattern of digits (**g, g’**); whereas those developing from more-proximal regions produce skeletal elements with proximal character, including one example of a miniaturised humerus (h*), radius (r*), and ulna (u*) (**h, h’**). Scale bar, 1 mm

Similar outcomes were obtained when 24 h ZPAEs were grafted to the anterior margin of host wing buds (Fig. 3e). Control ZPAE grafts produced mirror image duplications of the host digit pattern and contributed to the margin of the additional digit 3 (Fig. 3f; Supplementary Table 2). By contrast, cyclopamine-treated ZPAE grafts formed digit-like structures, often with multiple segments in patterns in which the host digits were duplicated (Fig. 3g; Supplementary Table 2). Grafts developing from more-proximal regions failed to duplicate host digits and gave rise to elements with proximal character, including, in one striking example, a miniaturised humerus together with an ulna and radius (Fig. 3h; Supplementary Table 2). Notably, similar results were reported in classic experiments in which chick wing bud mesoderm grafted to the chorioallantoic membrane regenerated an AER and produced segmented digit-like structures^19^. Together, these findings demonstrate that Shh signalling represses position-dependent skeletal differentiation within the ZPA.

### The Shh-p27*^Kip1^* pathway is absent in the mouse forelimb and hindlimb ZPA

Because p27*^Kip1^* prevents the chick wing ZPA from forming digits^9^, we investigated if it is expressed in the pentadactyl mouse forelimb and hindlimb, where the ZPA produces digits 4 and 5^1^. Using HCR in situ hybridisation we compared the distribution of *p27^Kip1^* with *Shh*. Whereas *p27^Kip1^* is specifically expressed in the chick wing and leg ZPA at HH24 and HH27 in a similar pattern to *Shh* ^8,20^ it is undetectable in the mouse forelimb and hindlimb at the equivalent stages of E10.5 and E11.5 (Fig. 4a-h). We also analysed the expression of the gene encoding Cyclin D2 which promotes G1-to-S-phase entry as part of an active kinase complex. In the chick wing and leg, *Cyclin D2* is restricted to a narrow posterior-distal domain that encompasses the ZPA at HH24 and HH27^8,20^ whereas in the mouse forelimb and hindlimb, it is broadly expressed throughout the distal mesoderm at E10.5 and E11.5 (Fig. 4i-p). These findings reveal that the cell cycle regulatory mechanism that prevents posterior digit development in the chick wing is absent in mouse forelimbs.

**Figure 4:**
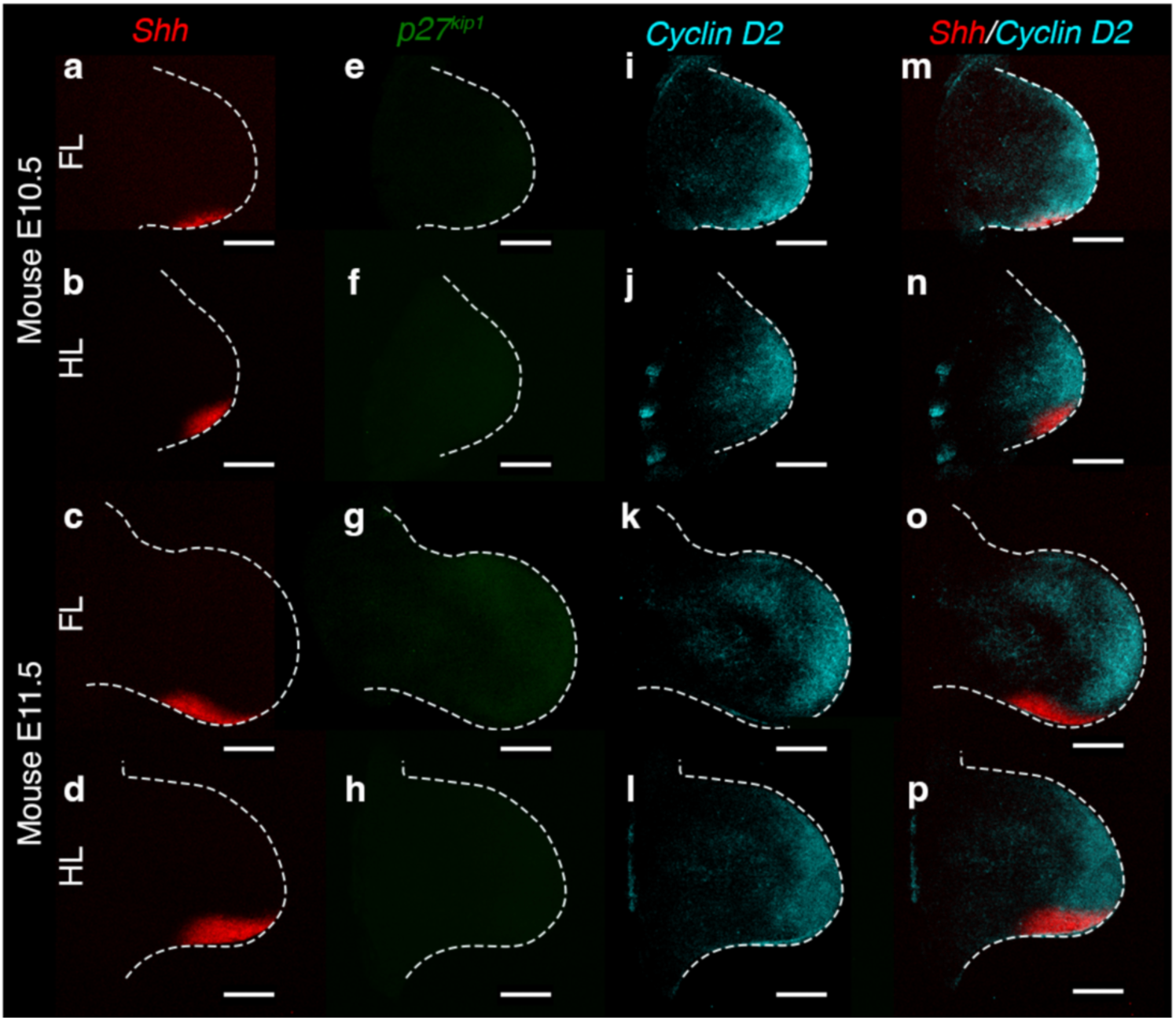
The Shh-p27^kip1^ pathway is absent in mouse limbs. **a–h**, Multiplex HCR-RNA FISH of embryonic day 10.5 (E10.5) mouse limbs. *Shh* (red) is detectable in E10.5 and E11.5 forelimbs (n = 5/5 and n = 8/8) and hindlimbs (n = 4/4 and n = 6/6) (**a-d**). *p27^Kip1^* expression (green) is undetectable in both E10.5 and E11.5 forelimbs (n = 8/8 and n = 12/12) and hindlimbs (n = 4/4 and n = 10/10) (**e-h**). *Cyclin D2* (cyan) is expressed broadly throughout the distal mesoderm of E10.5 and E11.5 forelimbs (n = 6/7 and n = 12/12) and hindlimbs (n = 4/4 and n = 9/10) (**i-l**). Minimal overlap between the broad distal *Cyclin D2* domain and the *Shh*-expressing ZPA of E10.5 and E11.5 forelimbs and hindlimbs (**m-p**). Scale bar, 500 µm.

### Transforming the chick leg into a mammalian-like digit pattern

The chick leg retains the stem amniote pattern of digit identities 1–2–3–4, defined by the ancestral phalangeal formula 2–3–4–5^3,5^, with digit 4 arising from the ZPA^2^ (Fig. 5a; progressive anterior-to-posterior specification of positional values over 16 h is also shown^2^). This conservation makes it an ideal system to study evolutionary changes in digit patterning since the divergence of mammals and birds. These considerations suggest that, in the stem amniote limb, digit 1-to-4 identities were specified as in the chick leg, and that digits 4 and 5 originated from the ZPA, as in mouse limbs (Fig. 5b)^1,5^.

**Figure 5:**
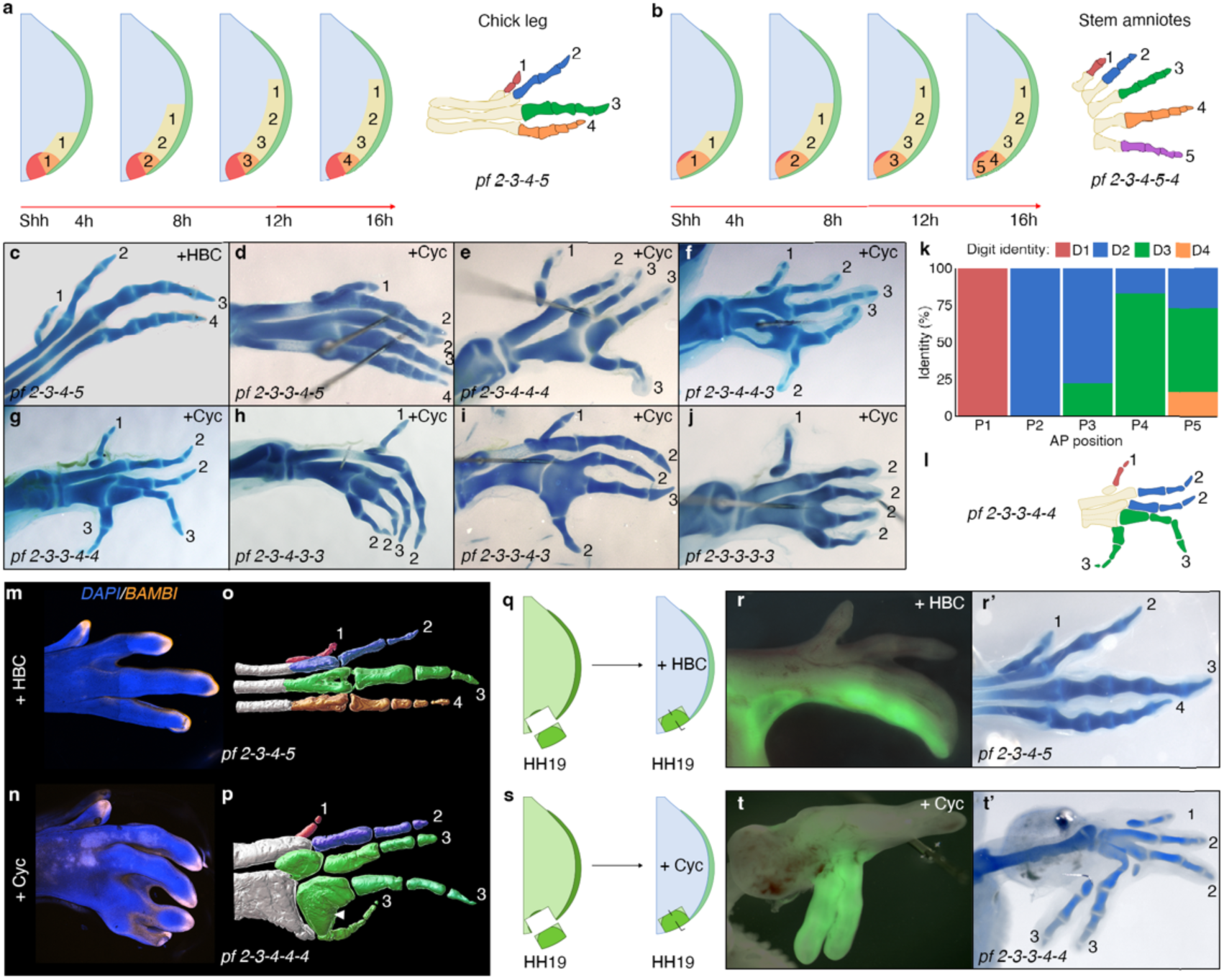
Transforming avian to mammalian modes of digit patterning. **a**, Specification of antero-posterior (AP) positional values in the chick leg. Positional values are specified over 16 h; transient Shh inhibition at 4-h intervals can generate a series of digit patterns (1-1, 1-2-2, 1-2-3-3, and 1-2-3-4)^2^. **b**, Ancestral stem amniote limb and predicted mechanism of AP specification. This model is based on the retention of the ancestral phalangeal formula (pf; 2-3-4-5) in the chick leg^3^ and the ability of the mouse limb ZPA to generate two digits^1^. **c–j**, Representative pentadactyl chick legs produced by cyclopamine application 8–10 h after *Shh* onset (**c**, HBC-treated control). Digit identities are assigned according to phalangeal formula: digit 1 (pf = 2), digit 2 (pf = 3), digit 3 (pf = 4), and digit 4 (pf = 5). **k**, Frequency distribution of digit identity relative to digit position in pentadactyl chick legs. **l**, Schematic of the most frequent pentadactyl pattern (1-2-2-3-3), with a pf of 2-3-3-4-4. **m**, **n**, HCR in situ hybridisation of *Bambi* mRNA. *Bambi* localises to the distal tips of all digits in control chick legs (m; n = 6/6) and pentadactyl legs (n; n = 9/9) indicating normal distal tip morphogenesis. **o**, **p**, Light-sheet imaging of normal and pentadactyl chick legs. Metatarsals and proximal phalanges are frequently fused in pentadactyl chick legs (arrowhead in p). **q, s**, Schematic of the GFP-expressing ZPA grafting procedure. **r, t** ZPA fate maps. In control legs, the ZPA contributes to a single posterior digit (n = 2/2) (**r**, **r’**). In pentadactyl legs, the ZPA contributes to two posterior digits (n = 2/2 legs with five digits) (**t, t’**)

We speculated that attenuating Shh signalling in the chick leg would cause ZPA overgrowth and additional digit formation as found in ZPAEs (Fig. 3). Within 4 h of exposing HH19/20 leg buds to cyclopamine, *Ptch1* expression was lost in the posterior part of the leg bud, demonstrating effective attenuation of Shh signalling during digit specification (Supplementary Fig. 4). Shh signalling inhibition also abolished *p27^Kip1^* expression within 24 h, correlating with ZPA over-proliferation by 10.2% at 48 h, as determined by flow cytometry (Supplementary Fig. 4). Additionally, the AER extended over the posterior part of the leg bud like in the mouse limb^8^, in association with de-repression of *Grem1* expression in adjacent posterior mesoderm by 48 h (Supplementary Fig. 4).

Precisely inhibiting Shh signalling in the chick leg at HH19/20 produced various pentadactyl patterns including the canonical mammalian configuration corresponding to digit identities 1–2–2–2–2 (pf, 2–3–3–3–3; Fig. 5c–j - Supplementary Table 3). Other embryos either lost posterior digits, or had a normal digit pattern, as previously reported^2^. Variation between phalangeal formulae in pentadactyl chick legs likely reflects subtle spatiotemporal differences in the timing of Shh inhibition (Fig. 5a). Positions two and three most often acquired digit 2 identity, whereas positions four and five typically acquired digit 3 and occasionally digit 4 identity (Fig. 5k). The most frequent outcome corresponded to identities 1–2–2–3–3 (pf, 2–3–3–4–4; Fig. 5l – Supplementary Table 3). Detailed skeletal analyses indicated that the digits were generally morphologically normal, with the distal-most phalanx tapering towards a *Bambi*-expressing tip, as observed in control limbs^21^ (Fig. 5m, n). However, the proximal phalanges were frequently fused, a feature better visualised by light-sheet imaging (Fig. 5o, p; white arrowhead in Fig. 5p; Supplementary Movies 1, 2).

To test whether the ZPA generates the fifth digit in cyclopamine-treated chick legs, we performed GFP-expressing tissue grafts at HH19 (Fig. 5 q, s). In controls, the ZPA produced the fourth digit^2^ (Fig. 5 r), whereas blocking Shh signalling caused it to produce both the fourth and fifth digits (Fig. 5t). Therefore, the precise removal of Shh signalling gives the chick leg ZPA a mammalian-like ability to generate two posterior digits in a pentadactyl pattern.

## Discussion

### Evolutionary determination of digit identity

Through precisely timed Shh pathway inhibition, we transformed the chick leg—which retains ancestral stem amniote digit identities—into mammalian-like pentadactyl patterns. The mechanism underlying this switch in developmental trajectory is summarised in our unifying model (Fig. 6a, c), suggesting that the digits of the mouse limb are specified through a comparable process, consistent with a transient requirement for Shh signalling^10^ (Fig. 6b). Our framework explains why pentadactyl chick legs resemble mouse hindlimbs morphologically at equivalent stages yet differ from normal chick legs (Supplementary Fig. 5).

**Figure 6:**
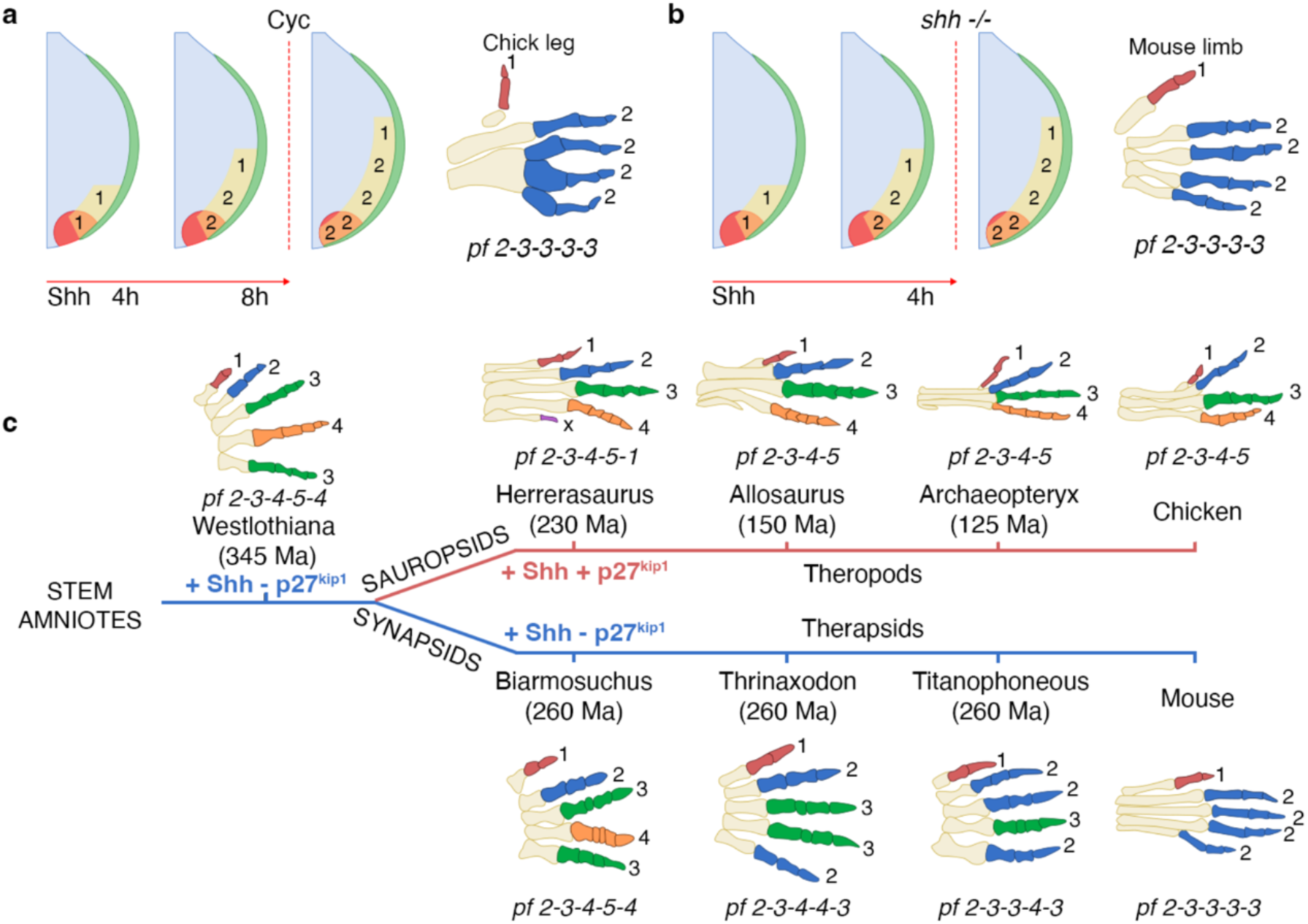
Evolution of avian and mammalian digit patterning. **a**, Schematic of the mechanism producing a 1-2-2-2-2 digit pattern (pf = 2-3-3-3-3) in the chick leg. Inhibition of Shh at the digit 2-to-3 promotion causes the ZPA to generate an additional digit 2. **b**, Extrapolation of the mechanism producing a 1-2-2-2-2 digit pattern in the chick leg to the mouse limb following the condition removal of Shh function (*shh^+/-^*)^10^. Differences in the duration of Shh required for a 1-2-2-2-2 digit pattern may reflect experimental approaches: genetic knockouts in mice^10^ versus abrupt signalling inhibition in chicks)^2^. Note, that the mouse limb pattern requires enforced cell survival following *Shh* deletion^10^. **c**, Evolution of digit patterning. In the stem amniote limb: digits specified by Shh as in chick leg; activation of the Shh–*p27^Kip^*^1^ pathway subsequently prevented digit 5 formation in the theropod/avian lineage—the same mechanism can explain digit 4 and 5 loss in the avian wing (Fig 1a). In mammals, digits specified by a brief pulse of Shh, producing a digit 1 and a uniform series of digit 2-like identities (pf = 2-3-3-3-3). Intermediate configurations are also shown that were present in non-mammalian therapsid ancestors (pf = 2-3-3-4-3; 2-3-4-4-3). Note, digits labelled by positional value, e.g., 1-2-2-2-2, rather than numerical order 1–5.

The spectrum of digit identities generated in pentadactyl chick legs mirrors those found in extinct therapsids (mammals and their immediate non-reptilian ancestors^6^ – Fig. 6c) and extant diapsid reptiles such as skinks^22^, geckos^23^, turtles^24^ and chameleons^25^, providing insight into evolutionary transitions across amniotes. Together, these comparisons suggest that temporal differences in Shh signalling are a principal determinant of digit identity, with longer durations progressively adding phalanges. However, the avian wing represents a derived exception to this rule: during the theropod dinosaur-to-bird transition the digits became developmentally truncated and lost terminal phalanges ^4,5,21^. Nevertheless, our ZPA explant grafting experiments reveal that the chick wing ZPA retains an ancestral capacity to produce multiply segmented digits, indicating that the underlying identity programme persists but is secondarily constrained.

### Evolutionary determination of digit number

Our explant grafting experiments indicate that ZPA size is a key determinant of digit number, potentially through a Turing-like reaction–diffusion mechanism^17,18^. Accordingly, when Shh signalling is attenuated, ZPAEs lose p27^Kip1^ expression, over-proliferate, self-organise local signalling interactions and form one or two digits after grafting into a host wing bud. The absence of the Shh–p27^Kip1^ pathway in mammals can therefore account for the pentadactyl pattern, in which two posterior digits originate from the ZPA^1^. The deployment of this pathway in archosaurs likely contributed to the reduction of digit number in wings (complete repression) and legs (partial repression) in modern birds, facilitating key evolutionary innovations such as flight and perching (Fig. 6c). Together, our results reveal two separable logics of Shh action: duration-dependent specification of digit identity and the p27^Kip1^-mediated control of digit number via regulation of ZPA growth.

---

**Supplementary Figure 1:**
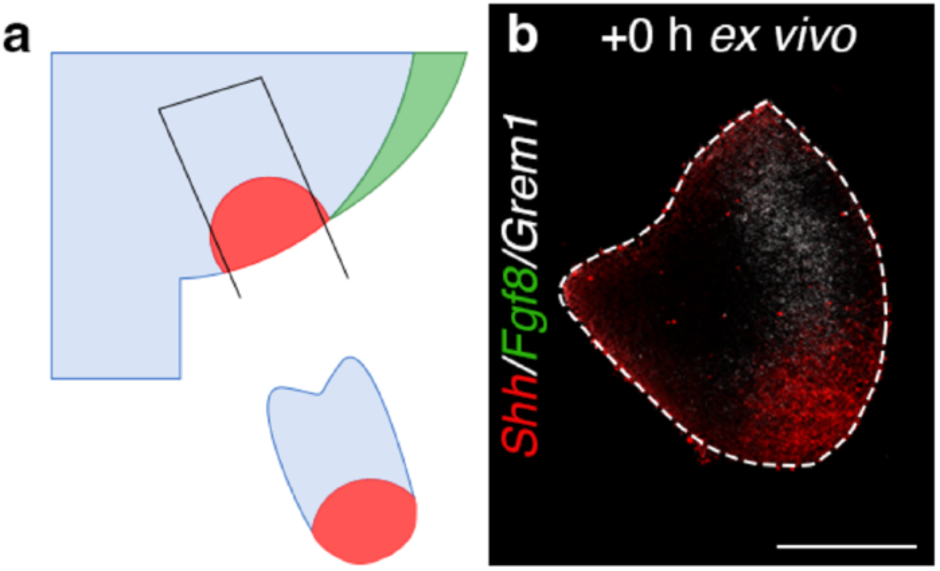
Validation of ZPA excision. **a**, Schematic illustrating the microsurgical dissection of the Zone of Polarising Activity (ZPA, red) from the surrounding mesenchyme (blue) and the Apical Ectodermal Ridge (AER, green). **b**, Multiplex HCR RNA-FISH of ZPA explants (ZPAEs) at 0 h. *Shh* (red) is expressed throughout the ZPA tissue, while the ZPAEs are devoid of *Fgf8*-expressing AER cells (green; n = 5/5). *Grem1* expression (white) is localised to the mesenchyme anterior to the *Shh*-expressing domain (n = 4/5). Scale bar, 250 µm.

**Supplementary Figure 2:**
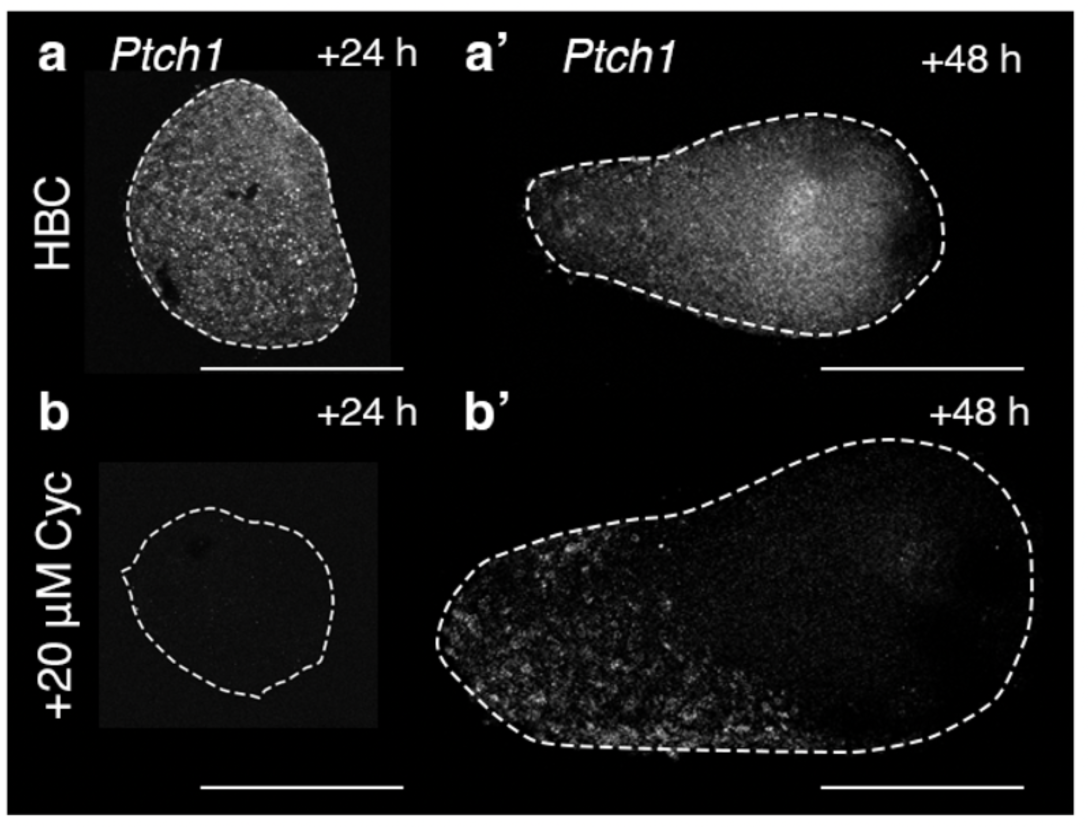
Verification of Shh pathway inhibition by cyclopamine. **a, a’**, ZPAEs treated with carrier-control (HBC) maintain robust expression of the Shh transcriptional target *Ptch1* at 24 h (**a**; n = 3/3) and 48 h (**a’**; n = 5/7), indicating sustained Shh signalling. **b, b’**, Application of 20 µM cyclopamine results in the loss of *Ptch1* expression by 24 h (**b**; n = 3/3), confirming effective pathway attenuation. Low basal levels of *Ptch1* are detectable by 48 h (**b’**; n = 5/6). Scale bar, 250 µm.

**Supplementary Figure 3:**
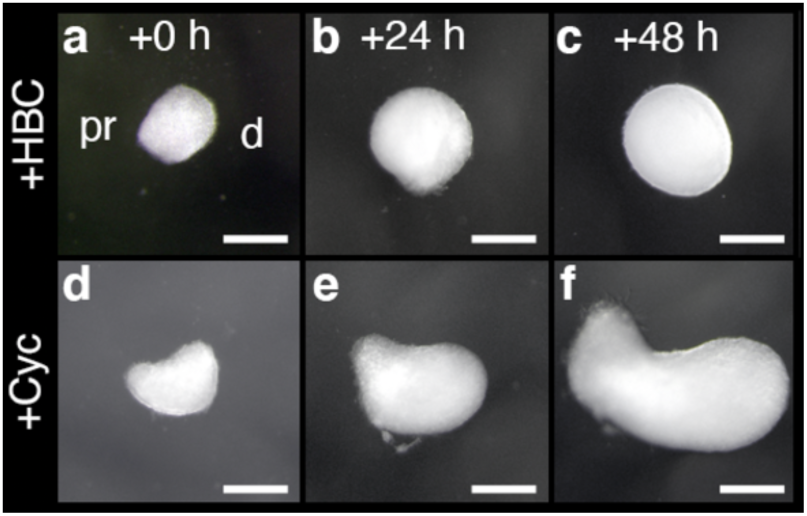
Shh inhibition causes ZPAE overgrowth. **a–c**, Brightfield images of control ZPAEs (HBC-treated) embedded in Matrigel. Explants exhibit minimal growth and retain a characteristic spheroid morphology over 48 h (n = 14/16). **d–f**, Cyclopamine-treated ZPAEs undergo pronounced expansion along the proximo–distal axis, resulting in an elongated shape by 48 h (n = 14/17). pr, proximal; d, distal. Scale bar, 250 µm.

**Supplementary Figure 4:**
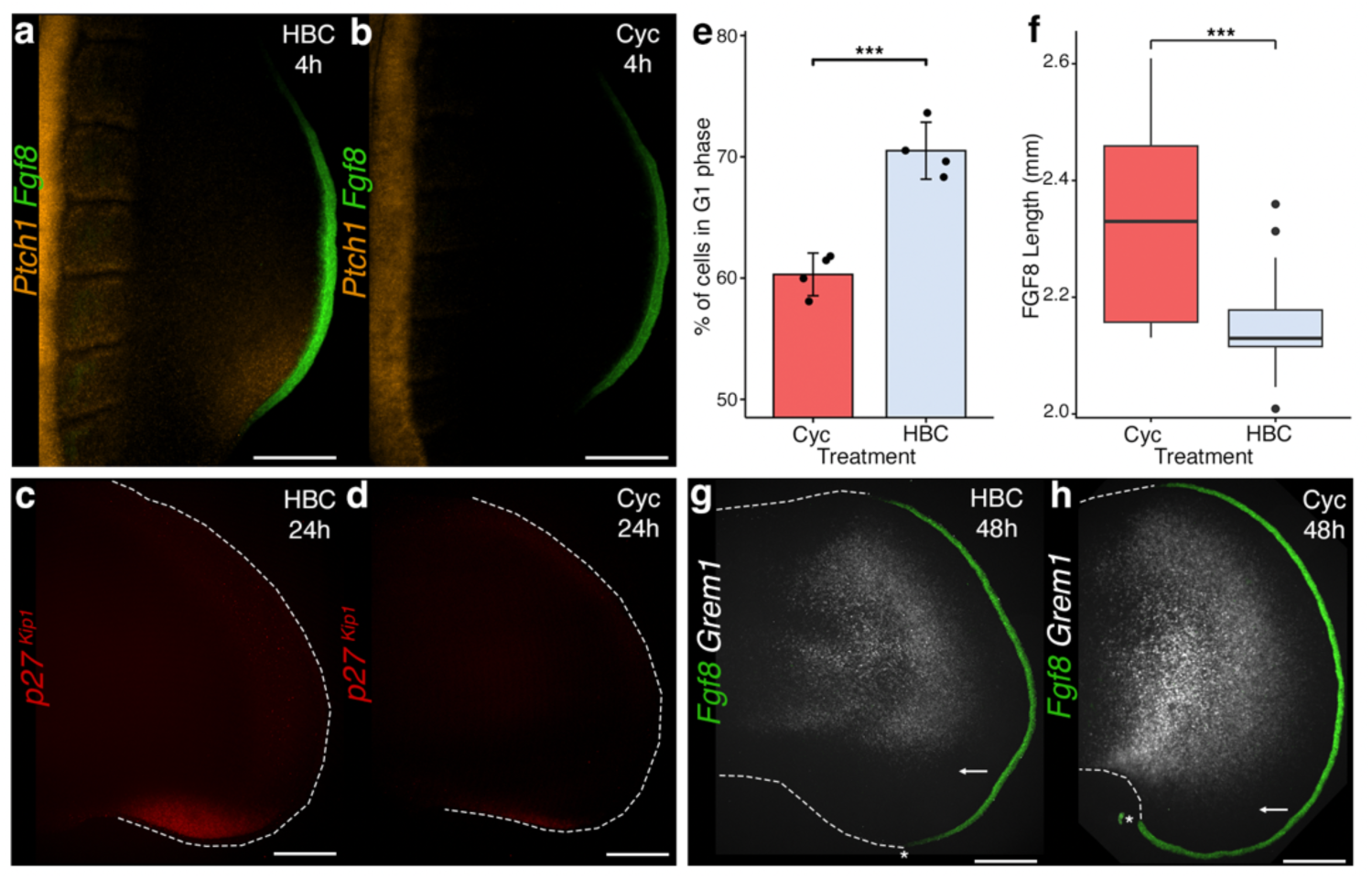
Consequences of Shh inhibition in ZPAEs. **a, b**, HCR RNA-FISH analysis of *Ptch1* (orange) and *Fgf8* (green) expression in HH19 embryos after 4 h of treatment with HBC (**a**; n = 3/4) or cyclopamine (**b**; n = 4/6). **c, d**, HCR RNA-FISH analysis of *p27^Kip1^* expression after 24 h. Cyclopamine treatment (**d**) abolishes *p27^Kip1^*expression compared to controls (**c**) (HBC n = 2/3; Cyc n = 3/3). **e**, Flow cytometric quantification of G1-phase cells at 48 h; cyclopamine treatment significantly reduces the G1 population, indicating accelerated G1-to-S progression (n = 4, 4; data represent mean ± s.d.; *P* = 0.0006, two-tailed unpaired t-test). **f**, Tukey style boxplot showing the length of the *Fgf8*-expressing AER at 48 h. Shh inhibition leads to a significant posterior expansion of the AER (n = 16 for HBC, n = 19 for cyc; *P* = 0.0003, two-tailed unpaired t-test). **g, h**, HCR RNA-FISH of *Grem1* (grey) and *Fgf8* (green) at 48 h; compared to controls (**g**) cyclopamine treatment (**h**) de-represses *Grem1* in posterior mesoderm near the extended AER (n = 3/3, white arrows). Scale bars 250µm.

**Supplementary Figure 5:**
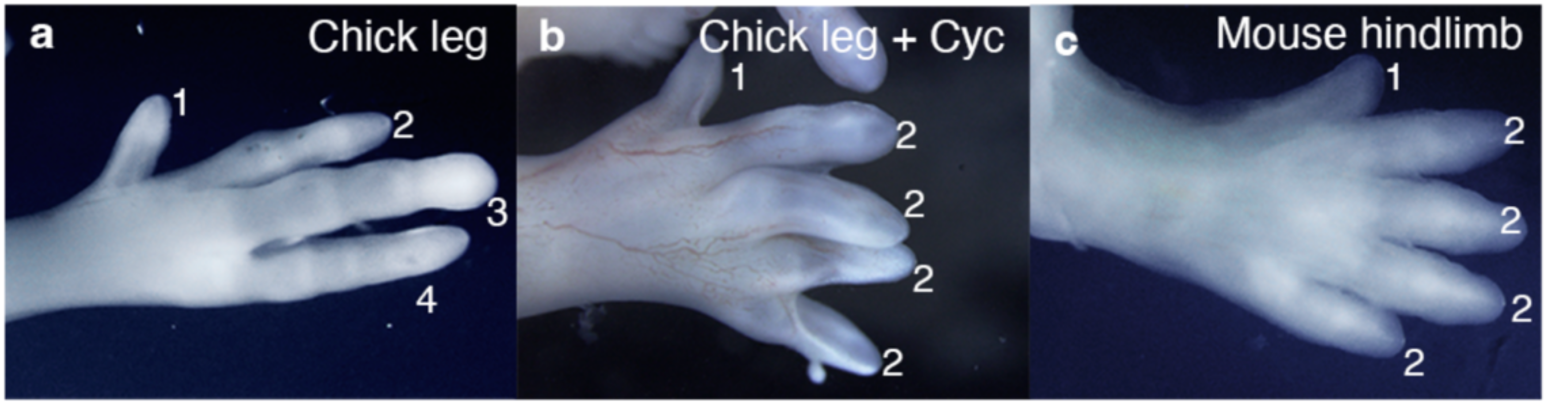
Morphological comparison of pentadactyl chick legs and mouse hindlimbs. **a**, Brightfield image of a chick leg at day 10 (D10) of incubation, showing the characteristic four-digit pattern. **b**, Chick leg treated with cyclopamine at HH18 results in the formation of an additional posterior digit, resembling the morphology of a mouse limb at an equivalent developmental stage. **c**, A mouse hindlimb at embryonic day 15 (E15) for comparison.

**Supplementary Table 1:** Number of skeletal structures arising from GFP*-*expressing ZPAE grafts. Grafting *GFP*-expressing ZPAEs, which had been incubated with cyclopamine for 24 h, into HH21/22 wing buds results in the formation of one or more ectopic skeletal structures (n = 3/4) compared to control grafts which did not form ectopic skeletal structures (n = 6/7). A single ectopic skeletal structure formed from ZPAEs grafted posteriorly in the autopod (n = 1/4), while 2 or more skeletal structures formed in the zeugopod (n = 1/4) and the autopod (n = 1/4). The identity of supernumerary structures is likely to reflect the area of the limb where the ZPAE integrated.

| POSTERIOR GRAFTS | HBC |  |  | Cyclopamine |  |  |
| --- | --- | --- | --- | --- | --- | --- |
| Location | Stylopod | Zeugopod | Autopod | Stylopod | Zeugopod | Autopod |
| 0 | 6/7 |  |  | 1/4 |  |  |
| 1 | - | - | 1/7 | - | - | 1/4 |
| 2 or more | - | - | - | - | 1/4 | 1/4 |

**Supplementary Table 2:**
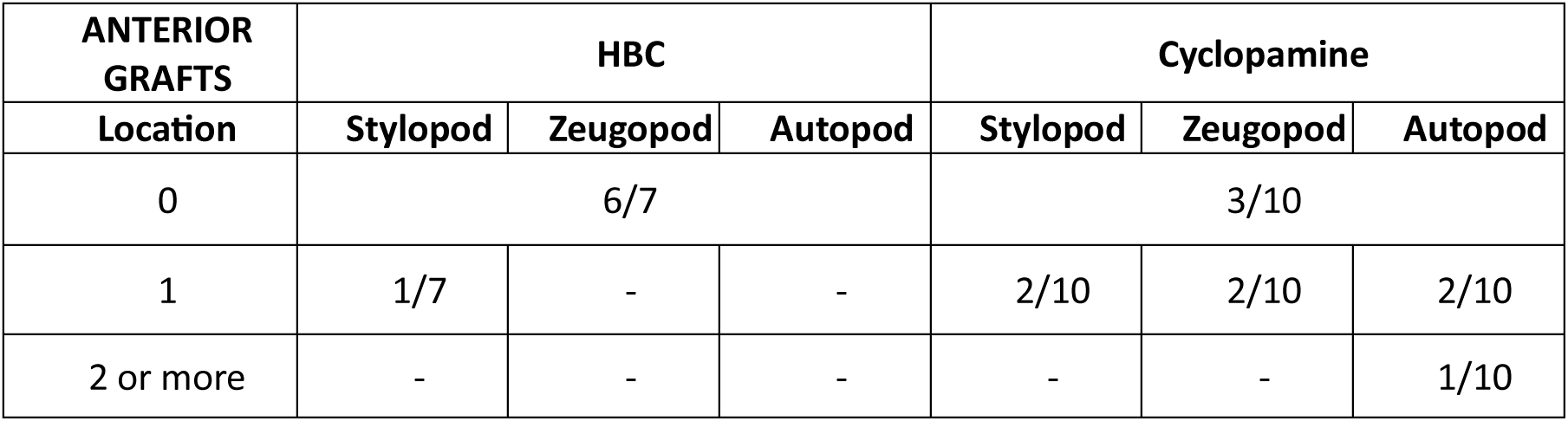
Number of skeletal structures arising from GFP*-*expressing ZPAE grafts. Grafting *GFP*-expressing ZPAEs, which had been incubated with cyclopamine for 24 h into HH21/22 wing buds, results in the formation of one or more ectopic skeletal structures (n = 7/10) compared to control grafts which did not form ectopic skeletal structures (n = 6/7). A single ectopic skeletal structure formed from a ZPAE in the stylopod (n = 2/10), the zeugopod (n = 2/10) and the autopod (n = 2/10). Additionally, two or more structures formed from a ZPAE in the autopod (n = 1/10). The identity of supernumerary structures is likely to reflect the area of the limb where the ZPAE integrated.

| ANTERIOR GRAFTS | HBC |  |  | Cyclopamine |  |  |
| --- | --- | --- | --- | --- | --- | --- |
| Location | Stylopod | Zeugopod | Autopod | Stylopod | Zeugopod | Autopod |
| 0 | 6/7 |  |  | 3/10 |  |  |
| 1 | 1/7 | - | - | 2/10 | 2/10 | 2/10 |
| 2 or more | - | - | - | - | - | 1/10 |

**Supplementary Table 3:** Alcian blue stained chick leg pentadactyl digit patterns. Targeted application of cyclopamine at HH19–20 produces pentadactyl digit patterns in the chick leg (n = 20) in ∼12% of cases. The low frequency of this phenotype likely reflects a narrow temporal window of sensitivity, which allows for sufficient posterior extension of the AER to support formation of an additional digit.

| Pattern | Count |
| --- | --- |
| 1-2-2-3-3 | 8 |
| 1-2-2-3-3 | 4 |
| 1-2-3-3-4 | 3 |
| 1-2-2-2-2 | 2 |
| 1-2-2-3-2 | 1 |
| 1-2-3-2-2 | 1 |
| 1-2-3-3-2 | 1 |

## Materials and Methods

### Chick and mouse husbandry

Wild type (Medeggs, Norfolk, UK) and GFP-expressing (Roslin Institute, Edinburgh, UK) Bovan Brown chicken eggs were incubated and staged according to Hamburger Hamilton^26^. Wild type mouse embryos (strain C57BL/6JH) were supplied by MRC Harwell (UK) at embryonic day (E) 10.5 and E11.5. All experiments were performed on embryos prior to the determination of sex, which is therefore not relevant to this study.

### Ex vivo polarising region explant (ZPAE) system

Polarising region (ZPA) tissue was dissected from HH20/21 wing buds in ice-cold PBS using a fine surgical knife under a Zeiss Stemi SV6 stereomicroscope. Beds of Growth Factor Reduced Matrigel (20µl; Corning) were prepared and polymerized in four-well plates for 45 min at 37°C. ZPAEs (4-5 per well) were placed on the Matrigel, overlaid with a second Matrigel layer, and cultured in CMRL medium supplemented with 10% FBS/1% Pen Strep/1% L-Glut in a humidified incubator with 5% CO_2_ at 37°C. Explants were recovered by replacing the culture medium with Cell Recovery Solution (Corning) and incubating on ice for 1h.

### Cyclopamine treatment

Cyclopamine (Sigma-C4116) was dissolved in hydroxybutyl-β-cyclodextrin (HBC) and added to the explant culture medium at a final concentration of 20µM at 0h. An equivalent concentration of HBC alone was used as a vehicle control. For embryo treatment, 5µl of 1mg/ml cyclopamine was pipetted onto HH18 chick embryos which were then allowed to develop for 6 days. An equivalent amount of HBC was pipetted onto HH18 chicks as a vehicle control.

### RNA-fluorescence in situ hybridisation with amplification by hybridisation chain reaction

Samples were fixed overnight in 4% PFA at 4°C, washed in PBS and dehydrated through a graded methanol series before storage in methanol at −20°C. Samples were rehydrated through a methanol-PBT series and treated with proteinase K for 3 min (explants) or 15-20 min (limb buds), followed by post-fixation in 4% PFA for 20 min. For limb buds, optional autofluorescence bleaching was performed by two washes in 3% H_2_O_2_/20mM NaOH/PBS solution for 45 minutes. Samples were washed sequentially in PBT, 5X SSCT and Probe Hybridisation Buffer (Molecular Instruments Inc.) before hybridisation with probes overnight at 37°C. All probes were custom generated (Molecular Instruments Inc.). Probes were prepared by diluting 5µl of 1µM probe in 500µl of Probe Hybridisation Buffer. Following hybridization, samples were washed with Probe Wash Buffer (Molecular Instruments Inc.), 5X SSCT and Amplification

Buffer (Molecular Instruments Inc.). Amplifier hairpins were heat-shocked at 95°C for 90s and snap-cooled at room temperature in the dark for 30min before incubation with samples in Amplification Buffer (Molecular Instruments Inc.) overnight at room temperature in the dark. Following this, samples were washed and stored in 5X SSCT before imaging. Fluorescent images of HCRs were obtained with either a Zeiss Apotome 2 microscope (10x objective) with Axiovision software (Zeiss) or a Nikon W1 Spinning Disk Confocal microscope (4X and 10X objective) with Nikon Elements Software. Images were processed with ImageJ v2.16.0 (FIJI) and Adobe Photoshop 2026.

### Colorimetric in situ hybridisation

Embryos/limb buds were fixed overnight in 4% PFA at 4°C, dehydrated in a methanol series, stored in methanol overnight at −20°C. Samples were rehydrated through a methanol-PBT series, washed in PBS, postfixed in 4% PFA at room temperature for 30 min and prehybridised in hybridisation buffer (50% formamide/50% 2X SSC) at 69°C for 2h. Samples were hybridised overnight at 69°C with antisense DIG-labelled mRNA probes diluted in prehybridisation buffer (1µg/1ml). Following hybridisation, samples were washed twice in hybridisation buffer, twice in a 1:1 mixture of hybridisation buffer-MAB buffer, and twice in MAB buffer then incubated in blocking buffer (2% blocking reagent/20% lamb serum in MAB buffer) for 2h at room temperature. Samples were incubated overnight at 4°C with anti-DIG antibody diluted in blocking buffer (1:2000) at 4°C, washed in MAB buffer overnight and developed in NTM buffer containing NBT/BCIP. mRNA expression patterns were visualised using a LeicaMZ16F microscope and LAS X 1.1.0.12420 imaging software.

### Explant size measurements

Explants were placed in a Petri dish containing 1xPBS and imaged using a Leica microscope and LASX 1.1.0.12420 imaging software. Explant surface areas were quantified using the Lasso tool with the Record Measurement Feature in Adobe Photoshop (2025).

### Flow cytometry for cell cycle analyses

Explants and polarising regions of stage-matched embryos were dissected in ice cold PBS under a Zeiss Stemi SV6 microscope using a fine surgical knife and pooled from replicate experiments (*n* = 10-14), before being trypsinised for 30 min into a single cell suspension. Cells were briefly washed twice in PBS and fixed in 70% ethanol overnight. Cells were washed in PBS and re-suspended and incubated for 20 min in PBS containing 0.1% Triton X-100, 50µg/ml propidium iodide and 50µg/ml^-1^ RNase A (Sigma). Single cells were analysed for DNA content with a Cytek Aurora flow cytometer using Flowjo or DeNovo FCSExpress. Single cells were gated using the forward scatter to determine which cells doublets and therefore excluded in the gate. Based on ploidy values, cells were assigned in G1, S or G2/M phases and this was expressed as a percentage of the total cell number (10,000 cells in each case).

### EdU labelling

Explants were incubated with 0.5 mM EdU in CMRL medium for 2 h at 37 °C, followed by fixation in 4% paraformaldehyde for 15 min. Samples were washed in 3% BSA in PBS at room temperature and permeabilized with 0.5% Triton X-100 in PBS for 20 min. EdU incorporation was detected using a click chemistry reaction (Azide Dye, Molecular Probes) for 1 h in the dark, according to the manufacturer’s instructions. Explants were subsequently washed in 3% BSA/PBS, counterstained with DAPI (1:1000 in 3% BSA/PBS) for 10 min and washed three times in 3% BSA/PBS prior to imaging. A Zeiss Apotome 2 microscope (10x objective) and Axiovision software (Zeiss) captured fluorescent images of EdU/DAPI labelled explants from which the percentage of EdU positive area was quantified using ImageJ v2.16.0 (FIJI) ^11^.

### Alcian blue cartilage staining

Day 10 embryos with viscera and head removed were fixed for 2 days in 90% ethanol. Embryos were then stained overnight in a 0.5% Alcian Blue 8GX (Sigma A5268) solution in 80%ethanol/20% acetic acid. Following staining, embryos were rehydrated in a graded ethanol series before being cleared in 1% KOH.

### ZPAE grafts

ZPAE of Roslin Green (Cytoplasmic GFP) embryos collected after 24h of culture in media containing either cyclopamine or HBC were grafted to wild-type HH21 embryos. A block of tissue (150 x 150 µm) was excised from the host wing bud at either the anterior or posterior edge using a sharpened tungsten needle. The ZPAE was then pinned into the excised space with an “L” shaped pin made from 25 µm platinum wire. The grafted embryos were then allowed to develop to Day 10 for cartilage staining.

### Light sheet fluorescence microscopy

Following RNA-FISH, samples were fixed overnight in 4% paraformaldehyde (PFA) at 4°C and dehydrated through a methanol series. After rehydration, samples were permeabilised overnight in PBST (PBS containing 0.5% Triton X-100) on a rocking platform. Samples were then stained with YO-PRO-1 iodide (1:300; Thermo Fisher Scientific) for 6 h. Following staining, samples were dehydrated through methanol and cleared according to the iDISCO+ solvent-based clearing protocol^27^. Imaging was performed using an UltraMicroscope Blaze (Miltenyi Biotec). Volumetric LSFM datasets were processed, segmented, and visualised using Imaris v11 (Oxford Instruments).

### Statistics & Reproducibility

All multiplexed hybridisation chain reaction (HCR), EdU labelling assays were performed on over 3 biological replicate explants (individual *n*-numbers are provided in respective figure legends and data is available in the Source Data file). All attempts at replication of these experiments were successful and we have included representative images with replicate information in the manuscript. To determine statistical significance in EdU labelling, two-tailed Student’s *t*-tests were used. To determine statistical significance of numbers of cells in different phases of the cell cycle (G1 vs. S, G2 and M) between pools of 8-12 explants in flow cytometry experiments, two-tailed Student’s *t*-tests were used. In all cases significantly different is taken as a *p*-value of less than 0.05. R (V4.6.0) and RStudio (2026.04.0+526) were used to construct graphs. No statistical method was used to predetermine sample size and no data were excluded from the analyses. The experiments were not randomised and the Investigators were not blinded to allocation during experiments and outcome assessment.

## Acknowledgements

We thank Cheryll Tickle and Marian Ros for critical reading and Graham Hickman (The Medical Technologies Innovation Facility (MTIF) at Nottingham Trent University) for help with light sheet imaging (BBSRC - BB/Z515760/1). We thank the Light Microscopy Facility at the University of Sheffield for confocal imaging (BBSRC – BB/M012522/1). MT is supported by the BBSRC (BB/Y0112615/1 and UKRI2946), M.P is supported by the Wellcome Trust (212247/Z/18/Z and 303188/Z/23/Z) and R.C is supported by the European Commission/Horizon Europe (101146024).

## Author contributions

S.W and S.S-P did the experimental work. R.C performed the light sheet microscopy. M.P contributed to developing the explant system. M.T designed the project and wrote the paper. All authors edited the manuscript.

## Materials and Correspondence

Should be addressed to

## Data availability

The authors declare that the] data supporting the findings of this study are available within the paper and its supplementary information files.

## Ethics declaration

The authors declare no competing interests.

